# Targeting glucocorticoid-induced CD20 activation in preclinical models of B-ALL

**DOI:** 10.64898/2025.12.16.694550

**Authors:** Jérémy Bigot, Mathieu Bouttier, Vincent Fregona, Clémence Rouzier, Manon Bayet, Sylvie Hebrard, Naïs Prade, Stéphanie Lagarde, Christine Didier, Laetitia Largeaud, François Vergez, Anne Quillet-Mary, Loïc Ysebaert, Cyril Broccardo, André Baruchel, Marion Strullu, Emmanuelle Clappier, Marlène Pasquet, Elodie Lainey, Eric Delabesse, Bastien Gerby

## Abstract

Pediatric B-cell acute lymphoblastic leukemia (B-ALL) is effectively controlled with contemporary multi-agent chemotherapy, resulting to 5-year survival rates above 90%. However, relapse occurs in 15-20% of patients due to minimal residual disease (MRD), characterized by the presence of persisting and resistant leukemic cells, and associated with a poor clinical outcome. Despite its prognostic relevance, the molecular features driving MRD are poorly characterized. In this study, we developed patient-derived xenograft (PDX) models from matched diagnosis and relapse B-ALL samples combined to chemotherapy to mimic MRD *in vivo*. Drug-tolerant leukemic cells were profiled using single-cell RNA sequencing and we identified a transcriptionally distinct MRD-like population enriched for cell-quiescence, inflammatory stress, and B-cell receptor pathway signatures. Strikingly, the B-lymphocyte surface antigen CD20, encoding by *MS4A1* gene, emerged as a consistent upregulated marker in MRD cells from PDXs and patients with diverse oncogenic subtypes. We further demonstrated that CD20 expression is induced by glucocorticoid exposure, creating a therapeutic opportunity where anti-CD20 monoclonal antibodies selectively eradicated MRD cells *in vivo*. Our data highlight CD20 not only as a biomarker but as an actionable vulnerability in B-ALL MRD, supporting clinical evaluation of anti-CD20 immunotherapy during induction treatment to kill drug-resistant cells and reduce relapse risk.

## Introduction

B-cell acute lymphoblastic leukemia (B-ALL) is an aggressive hematological malignancy classified into more than 20 distinct genetic subtypes. These oncogenic subgroups are established according to the identity of the first oncogenic event carried in the leukemic cells such as *ETV6::RUNX1*, *TCF3::PBX1*, *BCR::ABL1* translocations, *PAX5* alterations (fusion transcripts, *P80R* mutation), or chromosomal rearrangements involving *KMT2A* or *DUX4* (1). Current multi-agent chemotherapy is efficient at inducing long-term remission in child but is associated with severe side effects and undesirable consequences, including second malignant neoplasms (2). Indeed, while the 5-year survival rates now exceed 90%, the most common cause of treatment failure in pediatric B-ALL remains relapse that occurs in approximately 15–20% of patients (3). In addition, outcomes are particularly poor in adolescent and young adult (AYA) patients which are recognized as a unique population with specific disease biology (4). The prognosis is even worse in adult B-ALL because only 30% of adults achieve long-term disease-free survival (1). Although induction chemotherapies, combining glucocorticoids, asparaginase, anthracycline and vincristine, are efficient at reducing the tumor load by targeting proliferating and metabolically active leukemic cells, the minimal residual disease (MRD) arises in a significant proportion of patients and points to the presence of resistant cells that escape treatment (5).

The clinical utility of MRD assessment for the management of B-ALL patients is well-established. Indeed, early detection of MRD cells after induction chemotherapies has been consistently linked to important prognostic insights and remains the most powerful predictor of relapse as well as of overall survival in both pediatric and adult B-ALL (3,6,7). Although MRD monitoring has improved risk stratification, the mechanisms by which residual malignant cells escape chemotherapy remain to clarify and represent a key challenge for the development of targeted therapies. However, emerging studies demonstrated that cell plasticity, perturbation of lineage identity, cell quiescence and resistance appear tightly connected (8,9). These processes can be triggered by a non-cell-autonomous cue (10–12) or by an oncogene-induced molecular reprogramming, and should be considered for the eradication of MRD cells (13). Among the strategies used for MRD detection, multiparametric flow cytometry enables the discrimination of leukemic cells from normal B-cells, based on the expression of an aberrant phenotype (14). However, immunophenotypic modulation during the treatment can significantly impact MRD detection and leads to false negative results. While controverted, many markers used for MRD monitoring are affected by chemotherapies and glucocorticoid-induced expression modulation has been causally suspected (15–18). Therefore, therapy-induced activation of relevant pathways and/or surface markers can provide a window of therapeutic opportunity for killing drug-tolerant persister cells (19–21).

Here, we developed patient-derived xenografts (PDXs) to model MRD following induction chemotherapy in pediatric B-ALL. By using matched diagnostic and relapse samples from patients, and applying *in vivo* cytoreductive treatment, we aimed to characterize drug-tolerant leukemic cells that persist after therapy and to identify transcriptional programs and surface markers associated with resistance. Specifically, MRD cells from PDXs were profiled at the molecular and protein levels using single-cell RNA sequencing and FACS analysis. This strategy revealed a distinct sub-population with a unique transcriptional program enriched for cell-quiescence, inflammatory stress, and B-cell receptor pathway signatures. Among the genes consistently upregulated in MRD cells, we identified *MS4A1*, encoding the B-cell surface antigen CD20. Although targeting CD20 using immunotherapies, such as Rituximab and Obinutuzumab, has revolutionized the treatment of lymphomas (22) and has also shown clinical benefit in adult B-ALL (23,24), these therapies are not used for pediatric B-ALL. Here, we demonstrated that standard chemotherapies, particularly the use of glucocorticoids, lead to an increase in CD20 expression in MRD cells derived from both PDX models and pediatric patients, regardless of the oncogenic subtype. This mechanism opens a crucial therapeutic opportunity during induction therapy, allowing for the potent elimination of MRD cells using anti-CD20 monoclonal antibodies in childhood B-ALL.

## Results

### Experimental approach to detect MRD cells in human B-ALL

Based on the premise that chemotherapies are efficient at reducing the tumor load by targeting proliferating and metabolically active leukemic cells, we developed an experimental procedure to detect quiescent/resistant cells from matched diagnosis-relapse B-ALLs combining patient-derived xenograft (PDX) models and *in vivo* cytoreductive therapy (**Figure 1A**).

**Figure 1.**
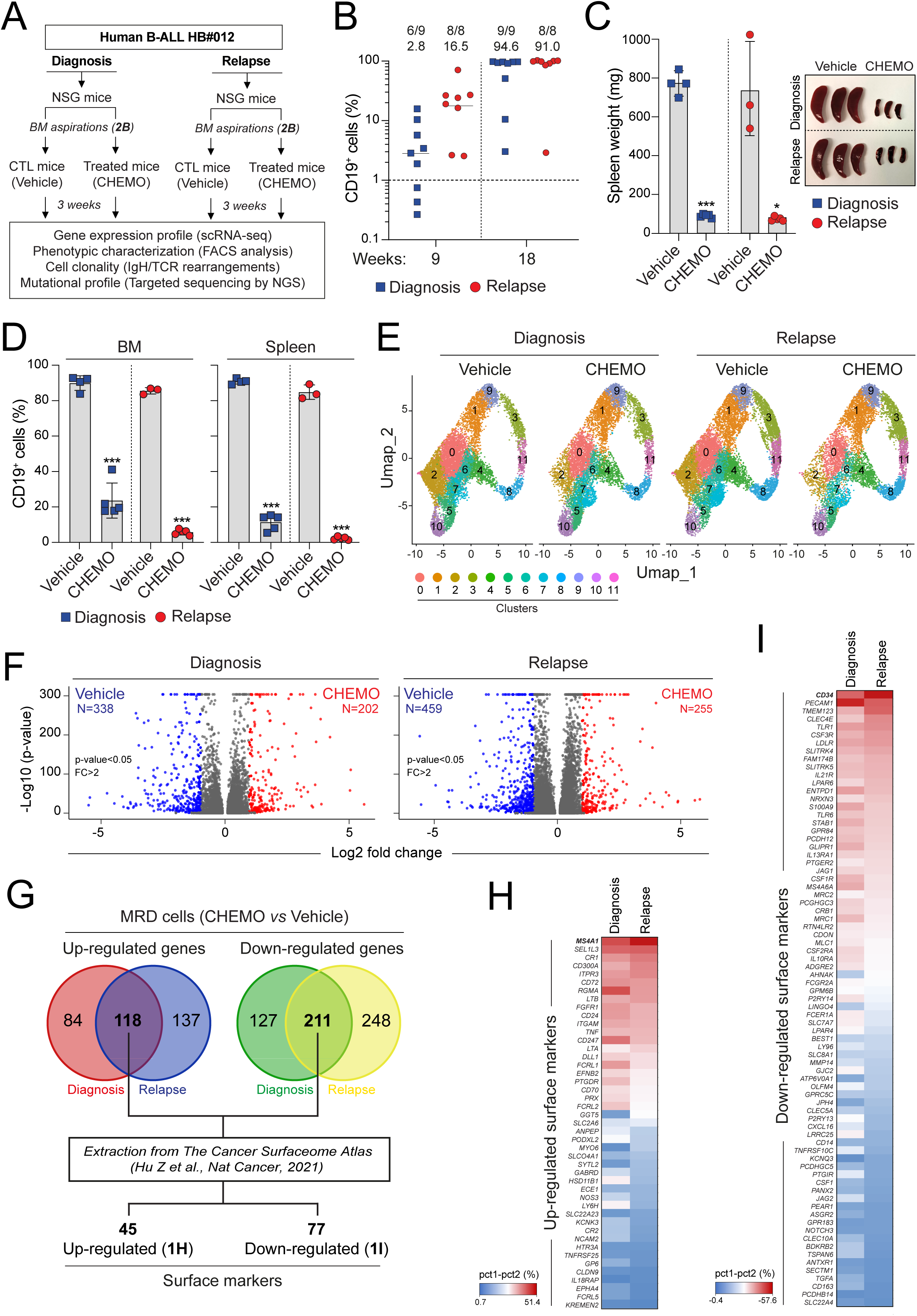
(**A**-**B**) Experimental procedure to study resistant cells from matched diagnosis-relapse B-ALLs. Diagnosis and relapse B-ALL leukemic blasts from patient HB#012 were transplanted into NSG mice (diagnosis n= 9, relapse n=8). Engrafted mice were randomly selected for treatment (CHEMO; diagnosis n=5 and relapse n=5) or not (vehicle; diagnosis n=4 and relapse n=3) with a chemotherapeutic cocktail (DEXA+VCR) during 3 weeks (A). The proportion of CD19^+^ blasts in BM aspiration was assessed 9 and 18 weeks after transplantation (B). (**C**) Spleen weight of each engrafted mouse was measured after treatment and picture of the representative spleens was shown. (**D**) Human B-ALL reconstitution (% of CD19^+^ B-ALL blasts) was monitored by FACS in the BM and the spleen after the treatment. (**E**) UMAP representation of single-cell RNA-seq data from treated (CHEMO) and untreated (vehicle) leukemic cells of the diagnosis and relapse HB#012 PDX models. Elven distinct transcriptional clusters of cells are distinguished by colors. (**F**) Volcano plot showing differentially expressed genes between vehicle and CHEMO leukemic from the diagnosis and relapse HB#012 PDXs. Up-regulated (red dots) and down-regulated (blue dots) genes are selected for an expression difference of >2-fold (p-value<0.05). (**G**) Venn diagrams showing overlap of common up-regulated (*left panel*) or down-regulated (*right panel*) genes in MRD cells after CHEMO from both diagnosis and relapse HB#012 PDXs. 47 up-regulated and 79 down-regulated genes encoding surface markers were identified by intersecting the gene lists with the human cancer surfaceome atlas (25). (**H**-**I**) Heatmaps showing the differential expression of the 47 up-regulated (H) and the 79 down-regulated (I) genes encoding surface markers after CHEMO at diagnosis and relapse, based on single-cell RNA-seq data (HB#012). The color scale represents the difference in the percentage of cells expressing each gene between treated (CHEMO) and untreated (vehicle) conditions (Δ% = pct₁ - pct₂ ; **Table S4**).

Leukemic blasts from a paired diagnosis (“*de novo*”) and relapse B-ALL patient with high hyperdiploidy (HB#012) were transplanted and expanded into immunodeficient NOD.*Cg-Prkdc^scid^Il2rg^tmWjl^*(refereed as NSG) mice (**Figures 1A**, **B**). Within 18 weeks, transplanted mice were highly and homogeneously engrafted, as assessed by bone marrow (BM) punctions (**Figure 1B**). Engrafted mice were then randomly selected to receive either Dexamethasone + Vincristine for 3 weeks (CHEMO) or vehicle alone (**Figure 1A**). B-ALL progression was prevented in treated mice as evidenced by a massive reduction of the leukemic burden in the spleen (**Figure 1C**). This drastic diminution of B-ALL cells *in vivo* was observed in both diagnosis and relapse samples, but was associated with the presence of persisting leukemic blasts in the BM and the spleen of recipient mice (**Figure 1D** and **Figures S1A, B**). These results suggest an equal sensitivity of leukemic cells from diagnosis and relapse upon CHEMO, and demonstrate our capability to detect residual resistant B-ALL cells using PDXs.

We also achieved the detection of resistant cells from a “*de novo*” *ETV6::RUNX1* B-ALL patient (HB#010) using the above protocol (8) (**Figures S1C-E**). To further characterize residual leukemic blasts from HB#010 *in vivo*, treated (CHEMO) and untreated (vehicle) cells from the BM were purified to perform gene expression profile using bulk RNA-sequencing (**Figures S1C-E**). Differential gene expression analysis revealed substantial transcriptional remodeling following treatment, with 687 genes up-regulated and 294 down-regulated (fold change > 3, FDR < 0.05) (**Figures S1F** and **Table S2**). To link our *in vivo* approach to the clinical situation, we compared their expression profile with a gene signature extracted from primary MRD cells purified by flow cytometry at the end of induction of five children (two *ETV6::RUNX1*, two hyperdiploid, one B-other) and two adult (*BCR::ABL1*) B-ALL patients (10). Our gene set enrichment analysis (GSEA) revealed a strong and significant enrichment of primary MRD signature in residual leukemic blasts from our HB#010 PDX model (**Figure S1G**). This transcriptional profile was associated with the dysregulation of the molecular pathways “TNFα signaling via NF-κB”, “inflammatory response”, “hypoxia” and “E2F targets” (**Figures S1H, I**), that have been previously associated with resistance and quiescence in B-ALL (8–10). Together, these molecular similarities indicate that our experimental approach combining PDXs and *in vivo* treatment mimics the clinical situation in B-ALL patients and allows for the detection of MRD cells.

### Molecular profiling identifies *MS4A1* as a gene candidate of MRD cells in PDX models

To further explore the transcriptional heterogeneity of untreated and MRD cells, we compared the gene expression profiles at the single-cell level (scRNA-seq) of treated (CHEMO) and untreated (vehicle) cells from the paired diagnosis-relapse HB#012 PDX models (**Figure 1A**). Unsupervised clustering approach using Uniform Manifold Approximation and Projection (UMAP) representation revealed twelve transcriptionally distinct clusters in each of the four conditions (**Figure 1E**), mainly distributed according to their status in the cell cycle (**Figure S2A**). The global analysis of the transcriptome identified 202 and 255 up-regulated genes in MRD cells from diagnosis and relapse, respectively (**Figure 1F** and **Table S3**). To find the most relevant cell-surface markers associated with the resistance of MRD cells, we overlapped them and we extracted from the Cancer Surfaceome Atlas (25) a list of 45 up-regulated genes encoding surface markers (**Figure 1G**, *left panel*). Among them, *MS4A1*, encoding the B-lymphocyte surface antigen CD20, was the gene for which the up-regulation was found in the most important proportion of cells between untreated and treated cells, from both diagnosis and relapse PDXs (**Figure 1H** and **Table S4**). Indeed, although *MS4A1* expression was mostly restricted to cluster 10 in untreated cells, *MS4A1* up-regulation was observed in all other clusters in MRD cells after CHEMO (**Figures 2A**, **B**, *upper panels*). Importantly, the overlap of the 45 up-regulated genes encoding surface markers in MRD cells from patient HB#012 (**Figure 1G**, *left panel*) with the up-regulated genes in MRD cells from patient HB#010 (**Figures S1F**) revealed *MS4A1* as a common gene, as well as *ITGAM* (*CD11b*) and *CD72*, that have been implicated for prognosis, treatment response monitoring and MRD detection in pediatric B-ALL (26–28) (**Figure S2B**).

**Figure 2.**
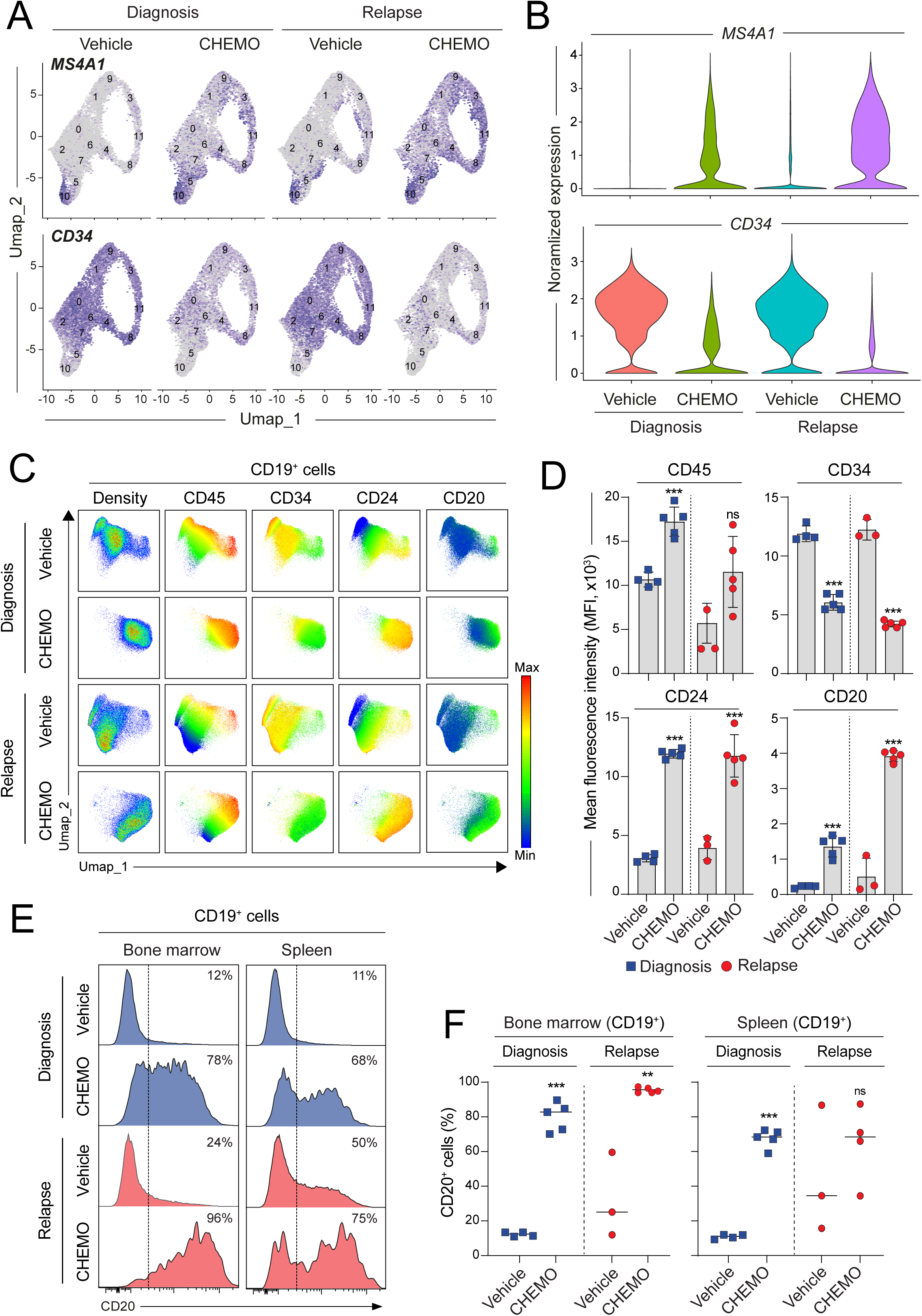
(**A**-**B**) Normalized expression levels of *MS4A1* (A, *upper panel*) and *CD34* (A, *lower panel*) shown on the UMAP representation of single-cell RNA-seq data from treated (CHEMO) and untreated (vehicle) leukemic cells of the diagnosis and relapse HB#012 PDX models. Each point represents a single cell, the color scale indicates the gene expression level and clusters are labeled (A). Violin plots showing the total distribution of *MS4A1* (B, *upper panel*) and *CD34* (B, *lower panel*) expression levels. (**C**-**D**) Phenotypic characterization by UMAP representation of human CD19⁺ B-ALL blasts from the BM of diagnosis and relapse HB#012 PDXs treated (CHEMO) or not (Vehicle). Visualization of the cell density (C, *left panels*) associated with the expression level (C, *right panels*) and the mean fluorescence intensity (D) of CD45, CD34, CD24 and CD20 was shown. (**E**-**F**) Representative FACS analysis (E) and quantification (F) of CD20 expression in CD19⁺ B-ALL blasts from the BM and the spleen of diagnosis and relapse HB#012 PDXs treated or not with CHEMO.

Next, we performed the same analysis with down-regulated genes in MRD cells from diagnosis and relapse PDXs (**Figure 1F** and **Table S3**). A list of 77 genes encoding cell-surface markers was established (**Figure 1G**, *right panel*) and we identified *CD34* as the most relevant candidate for which the down-regulation was associated with the resistance of MRD cells (**Figure 1I** and **Figures 2A**, **B**, *lower panels*). Interestingly, cluster 10 was the only population within untreated cells, from both diagnosis and relapse PDXs, that expressed *MS4A1* and simultaneously repressed *CD34* (**Figure 2A**). Therefore, we further investigated the molecular signatures of cluster 10 from the leukemic bulk and we observed a strong enrichment of primary MRD signature (10) as well as of cell-quiescence signature (29,30) (**Figure S2C**). Furthermore, our gene set variation analysis revealed BCR signaling as the most enriched molecular pathway in cluster 10 (**Figure S2D**). Finally, we confirmed our findings by comparing the enrichment score of these three signatures between *MS4A1* positive and negative cells from the scRNA-seq data (**Figure S2E**). BCR being reported to interact directly with CD20 (31,32), our results suggest the presence within the tumor bulk, of a minor cell population exhibiting a mature immunophenotype CD34^-^CD20^+^.

### Chemoresistance is associated with CD20 up-regulation in PDX models

To confirm our findings at the protein level, we performed a multiparametric staining by FACS covering the steps of human B-cell differentiation (**Figure S2F** and **Table S5**). Unsupervised clustering revealed a phenotypic evolution of MRD cells as compared to untreated cells, characterized by the down-regulation of CD34 and the up-regulation of CD45, CD24 and CD20 (**Figures 2C**, **D**), while the expression of CD10 and CD38 markers remained stable (**Figure S2G**). This phenotypic progression of residual cells toward B-cell maturation observed in diagnosis PDXs was even more pronounced in relapse PDXs (**Figures 2C**, **D**), corroborating our molecular observations (**Figures 2A, B**). Indeed, CD20^+^ cells were significantly enriched in MRD cells from the BM and the spleen of diagnosis PDXs, and this enrichment was particularly strong in relapse PDXs after CHEMO (**Figures 2E, F**). In addition, we confirmed the presence of CD34^-^CD20^+^ population within the untreated bulk (**Figures 2C**), probably corresponding to the cells from cluster 10 of the scRNA-seq data (**Figure 2A**). Finally, we compared the cell clonality of untreated and MRD cells by analysing their IgH/TCR rearrangements and by exploring their mutational profiles by targeting sequencing of genes recurrently mutated in B-ALL patients (**Figure 1A** and **Tables S6**, **7**). The data revealed a same and unique clone exhibiting the TCRδ V2-D3 gene rearrangement associated with the presence of KRAS^G12R^ mutation between diagnosis and relapse samples from the patient, and from untreated or treated PDXs with CHEMO (**Figure S3A**). We also controlled by cell purification, that the minor population of CD20^+^ cells derived from the same clone with similar variation allele frequencies (VAF) of TCRδ V2-D3 rearrangement and of KRAS^G12R^ mutation (**Figure S3B**). These results strongly suggest that the transcriptional and phenotypic modifications observed in MRD cells from our HB#012 PDX models are due neither to genetic heterogeneity of leukemic cells nor to clonal selection and/or evolution processes after CHEMO.

The above results indicate that CD20 up-regulation is associated with chemoresistance in PDX models. To strengthen this idea, we controlled that CD20^+^ cells were enriched in MRD cells from the BM and the spleen of HB#010 PDX models (**Figure S3C**) and we repeated our *in vivo* experimental approach with another matched diagnosis-relapse B-ALL patient carrying *ETV6::RUNX1* rearrangement (HB#020) (**Figure 3A**). We confirmed that the treatment with CHEMO for 3 weeks efficiently prevents B-ALL progression in the spleen and in the BM of both diagnosis and relapse PDXs (**Figures S3D, E**), and is associated with the presence of persisting leukemic blasts exhibiting the immunophenotype CD45^high^CD34^low^CD20^+^ (**Figures 3B**, **C** and **Figure S3F**). To further investigate the BCR pathway activation in CD20^+^ leukemic cells, we used a phosphoflow cytometry approach after chemical *ex vivo* stimulation. BCR signaling response was assessed by mimicking its downstream activation with hydrogen peroxide (H2O2), a well-known inducer of PLCγ2 phosphorylation (33,34). As expected, H2O2 induced the phosphorylation of PLCγ2 more efficiently in purified CD20^+^ than in CD20^-^leukemic cells (**Figure S3G**). Together, our results indicate a phenotypic heterogeneity of B-ALL leukemic blasts, associated with the presence of a minor population of quiescent CD20^+^ cells, exhibiting a functional BCR and that are resistant to chemotherapy.

**Figure 3.**
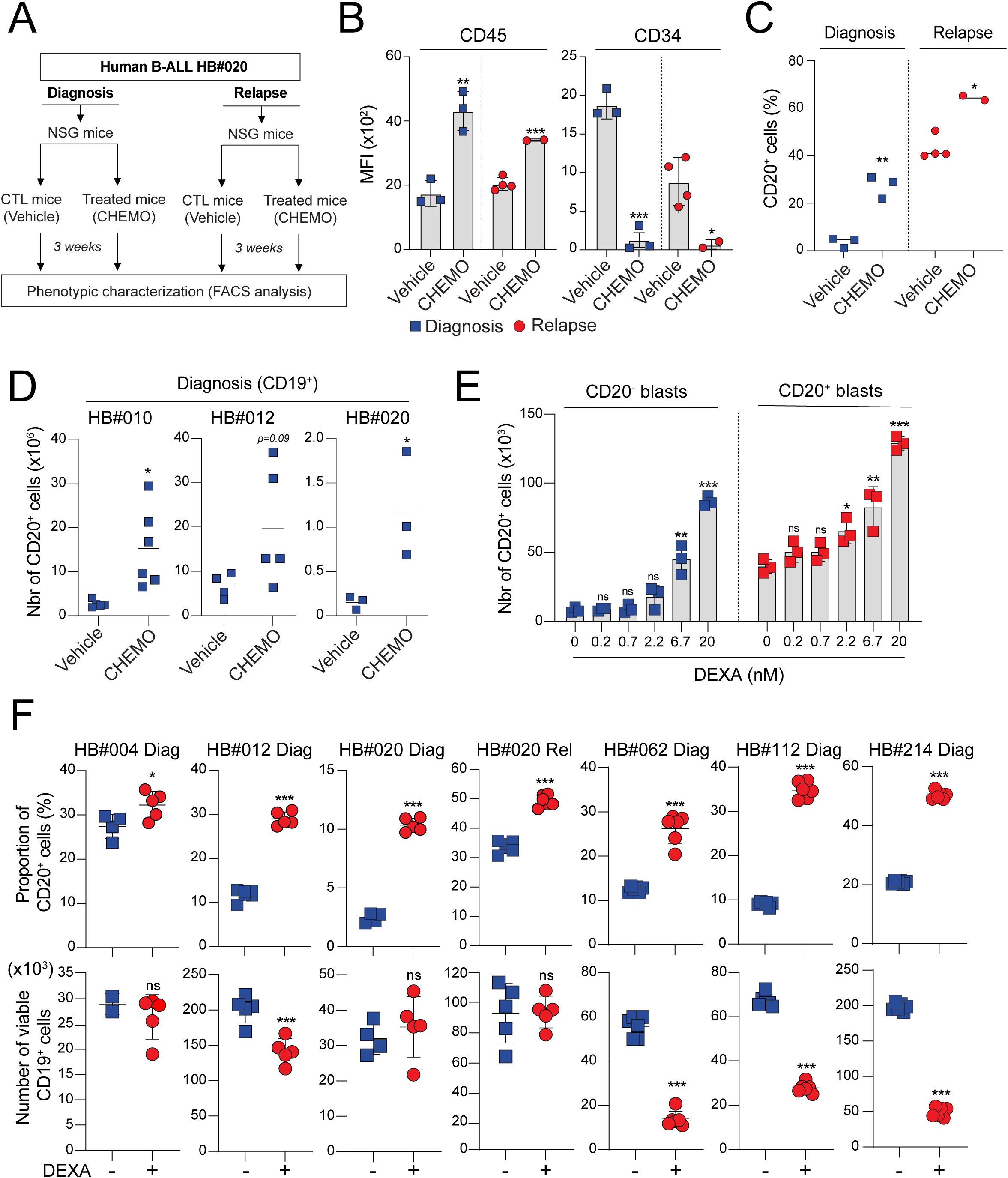
(**A**-**C**) Experimental procedure to explore the immunophenotype of chemo-resistant B-ALL cells. Diagnosis and relapse B-ALL leukemic blasts from patient HB#020 were transplanted into NSG mice. Engrafted mice were randomly selected for treatment (CHEMO; diagnosis n=3 and relapse n=5) or not (vehicle; diagnosis n=3 and relapse n=5) with a chemotherapeutic cocktail (DEXA+VCR) during 3 weeks (A). Human B-ALL reconstitution (% of CD19^+^ B-ALL blasts) was monitored by FACS in the BM and the spleen after the treatment (**Figure S3E**). MFI of CD45 and CD34 in engrafted CD19^+^ B-ALL blasts in the BM of recipient mice was shown (B) and CD20 expression was quantified (C). (**D**) Absolute number of CD20⁺ B-ALL blasts (CD19⁺) from the BM of diagnosis HB#010, HB#012 and HB#020 PDXs treated or not with CHEMO. (**E**) CD20^+^ (blue) and CD20^-^ (red) leukemic cells from relapse HB#012 PDXs were purified and treated *in vitro* on MS5 stromal cells with a dose-response of DEXA (0 to 20nM, n=3) for 48h. Absolute number of CD20⁺ B-ALL blasts (CD19⁺) was then calculated for each condition. (**F**) CD20 expression on CD19^+^ B-ALL blasts from HB#004, HB#012, HB#020, HB#062, HB#112 and HB#214 PDXs was quantified after 48h of co-cultured on MS5 stromal cells in presence or not of 60nM DEXA (*upper panel*). Absolute number of CD19^+^ B-ALL blasts was measured under the same conditions (*lower panel*).

### CD20 is a plastic marker modulated by dexamethasone

Despite a significant reduction of leukemic cells, we noticed that the treatment with CHEMO led to an increase not only in the proportion but also in the absolute number of CD20^+^ blasts in the BM of HB#010, HB#012 and HB#020 PDXs (**Figure 3D**). This observation supports the notion that CHEMO could not only select pre-existing CD20^+^ blasts, but also induce CD20 expression at the surface of leukemic cells. This dual effect may explain the widespread *MS4A1* up-regulation observed across all the clusters after CHEMO in our scRNA-seq data (**Figure 2A**, *upper panel*). Thus, we addressed the question whether CD20 expression is modulated by dexamethasone (DEXA) *in vitro*. A dose-response experiment of DEXA on leukemic cells from HB#012 PDXs, which initially displayed 12% of CD20^+^ cells (**Figure 2E**), was associated with an important up-regulation of CD20 in a dose-dependent manner, reaching ∼35% of CD20^+^ cells with 60nM DEXA (**Figures S3H**). To confirm this inducibility of CD20 expression, purified CD20^+^ and CD20^-^ leukemic blasts were treated in dose-response with DEXA (0 to 20nM), a range chosen to avoid cytotoxicity. From the CD20^-^ fraction, we observed a clear dose-dependent increase in both frequency and absolute number of CD20^+^ cells (**Figure 3E** and **Figure S3I**, *left panel*), while total CD19⁺ cell counts remained stable (**Figure S3I**, *right panel*). This confirms that DEXA actively induces CD20 expression in cells that were initially negative. Interestingly, the proportion of CD20 from purified CD20^+^ fraction dropped to 19% after the co-culture, but increased to 55% when treated with 20 nM of DEXA (**Figure 3E** and **Figure S3I**, *left panel*), indicating that CD20 expression is plastic and subject to steroid-driven modulation. To assess this phenomenon across other B-ALL subtypes, we then examined six additional B-ALL PDX models, including three with the oncogenic subtype PAX5^P80R^ (HB#062, HB#112, HB#214) (**Table S3**). We showed that treatment with 60nM DEXA systematically increased the percentage of CD20^+^ cells (**Figure 3F**, *upper panel*), regardless of its impact on cell viability (**Figure 3F**, *lower panel*). Together, our results demonstrate that chemoresistance is associated with CD20 expression at the surface of MRD cells in PDX models, and that DEXA induces CD20 up-modulation at the surface of viable leukemic blasts *in vitro*.

Finally, to address the functional contribution of CD20^+^ and CD20^-^ cells in leukemia progression, the kinetic of CD20 expression during B-ALL progression in PDXs was monitored (**Figure 4A**). Strikingly, CD20^+^ population was highly enriched in short-term engrafted cells, while remained progressively diluted in the leukemic bulk after medium- and long-term reconstitution (**Figure 4B**). Indeed, we found that CD20 expression was inversely correlated with the tumor load (**Figure 4C**), supporting the view that CD20^+^ and CD20^-^ fractions could contribute to leukemia development with different kinetics. To directly address this question, equal number of CD20^-^ and CD20^+^ blasts were purified and transplanted in NSG mice (**Figures 4D-F**). While both fractions were capable to engraft, we observed that mice receiving CD20⁺ subset displayed slower kinetics of leukemia progression compared to those receiving CD20^-^ cells (**Figure 4E**). Notably, at early time post transplantation, a substantial proportion of CD20^+^ emerged in mice initially transplanted with the CD20^-^ fraction, supporting a phenotypic conversion *in vivo*. However, CD20⁺ cells became progressively diluted within expanding leukemic cells over time (**Figure 4F**). Collectively, our results confirm the plastic state of CD20 *in vivo* and reinforces the notion that quiescent/resistant CD20^+^ cells exhibit a long-term leukemia initiating activity, mimicking in some aspect relapse-initiating cells from B-ALL patients.

**Figure 4.**
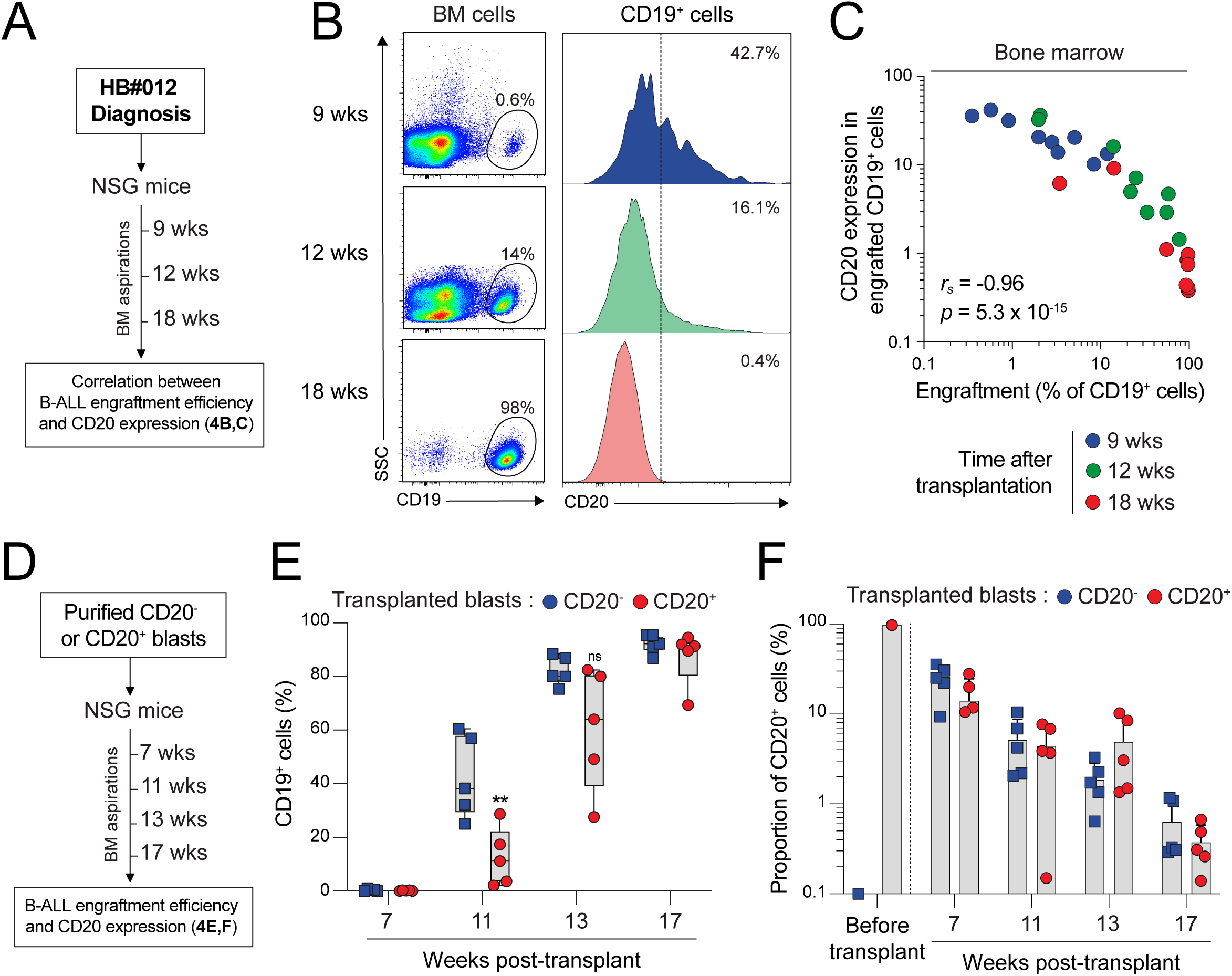
(**A-C**) Kinetic of engraftment 9, 12, and 18 weeks after transplantation of leukemic blasts from HB#012 patient at diagnosis (A). Representative FACS analysis of B-ALL reconstitution (B, *left panel*, % of CD19^+^ B-ALL blasts) in the BM and of CD20 expression on engrafted cells (B, *right panel*) for each time point. Negative correlation between engraftment efficiency (% CD19⁺ cells) in the BM and CD20 expression on leukemic cells (C). (**D-F**) Equal number of CD20^+^ (blue) and CD20^-^ (red) leukemic blasts (1×10^5^ CD19⁺) from relapse HB#012 PDXs were purified and transplanted into NSG mice (D, CD20^+^, n=5 and CD20^-^, n=5). Kinetic of B-ALL reconstitution (E, % of CD19^+^ B-ALL blasts) and CD20 expression on engrafted cells (F) were monitored by FACS in the BM at indicated time points post-transplantation.

### CD20 is a marker of MRD cells in B-ALL patients during induction treatment

To validate our findings in a clinically relevant context, we analyzed a cohort of 340 pediatric B-ALL patients receiving an induction therapy combining vincristine, PEG-asparaginase, and dexamethasone, supplemented with daunorubicin for NCI-high risk patients (<1Y or >=10y and/or >=50G/L). From this cohort, a subgroup of 45 patients (13.2%) with MRD>0.3% at day 15 of induction therapy was selected to evaluate the phenotypic evolution of blasts by flow cytometry during treatment. The analysis revealed a significant down-regulation of CD34 and up-regulation of CD45 and CD20, while CD19 expression remained stable (**Figure 5A** and **Table S8**), strongly supporting our results in MRD cells from the PDX models (**Figures 2C-F** and **Figures 3B, C**). This was associated with a global and massive increase of CD20⁺ blasts in MRD cells at day 15 after treatment compared to leukemic cells at diagnosis (**Figure 5B** and **Table S8**). We therefore established five categories of patients (groups I to V), according to the evolution of CD20 expression during treatment (**Figures 5C** and **Figure S4A**). Strikingly, more than half of patients (23/45), corresponding to groups II and III, displayed a marked induction of CD20 at day 15 (**Figure 5C**). In particular, patients from groups II (18/45), who were considered negative for CD20 (<20% of CD20^+^ blasts) at diagnosis, became positive (>20% of CD20^+^ blasts) at MRD (**Figure 5C** and **Figure S4A**). This patient category demonstrates how therapy-driven immunophenotypic modulation can alter the MRD monitoring by flow cytometry during early response assessment. In addition, while initially considered CD20 positive at diagnosis, patients from groups III (5/45) exhibited a significant CD20 up-modulation at MRD (>50% of CD20^+^ blasts) (**Figure 5C** and **Figure S4A**). Finally, CD20 activation in MRD cells was observed in diverse oncogenic subtypes of B-ALL (*ETV6::RUNX1,* high hyperdiploid, *IGH::DUX4* and B-other patients). This suggests that CD20 up-modulation during chemotherapy is a widespread and oncogene-independent feature of MRD cells in patients (**Figure 5D** and **Figure S4B**). Together, our findings corroborate the phenotypic plasticity of leukemic cells under therapeutic pressure and establish CD20 as a robust and clinically relevant marker of drug-tolerant blasts during the early phase of treatment.

**Figure 5.**
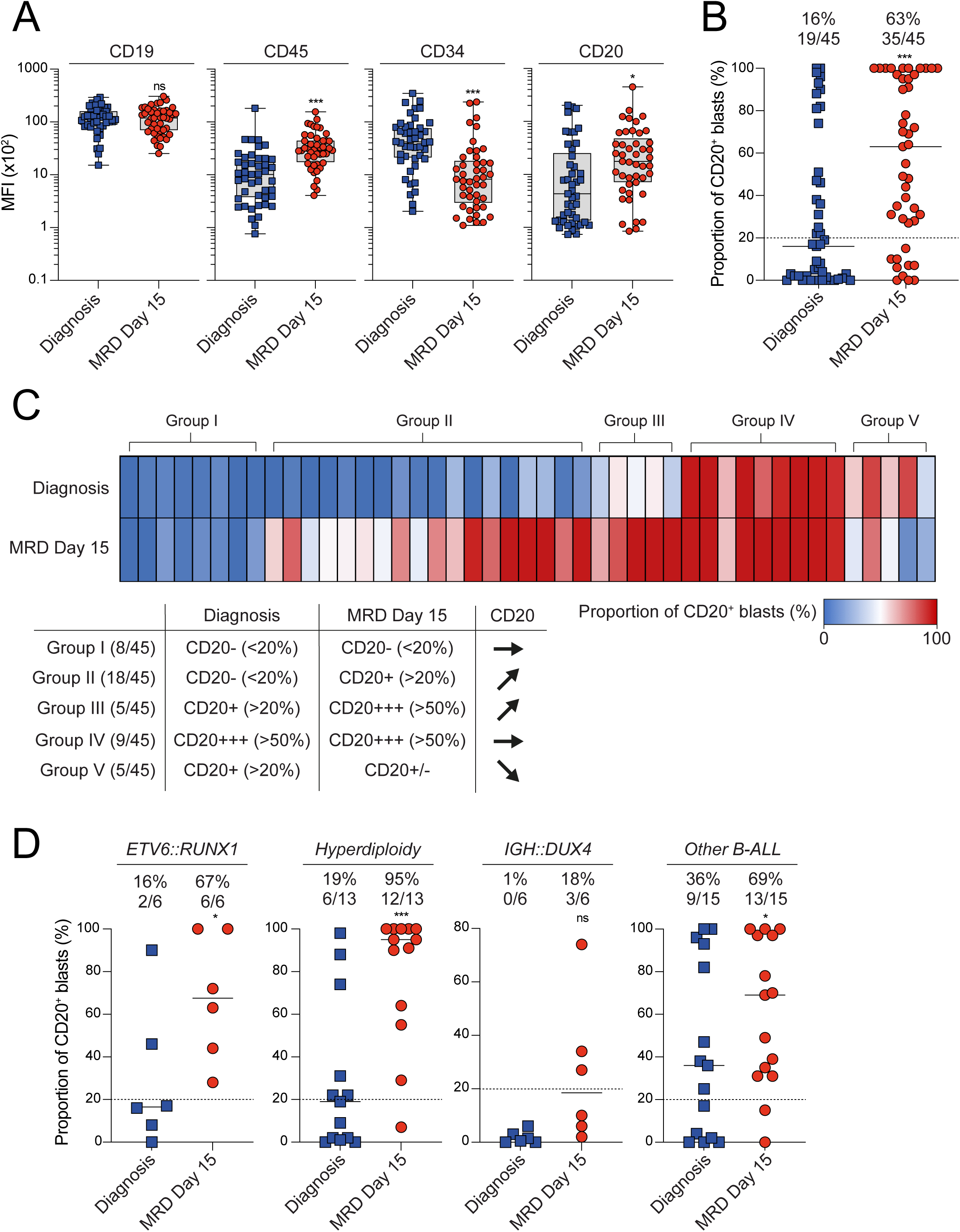
(**A**) CD19, CD45, CD34 and CD20 mean fluorescence intensity (MFI) analysed by FACS of leukemic blasts at diagnosis and at day 15 of induction therapy (MRD Day15) in 45 pediatric B-ALL patients (MRD >0.3%). (**B**) The proportion of CD20⁺ cells among CD19⁺ blasts at diagnosis and at day 15 in the cohort was analysed. Median of CD20 expression and percentage of positive patients (>20%) were indicated. (**C**) Heatmap representation of CD20⁺ cell frequency at diagnosis and MRD (*upper panel*), with patients grouped into five categories (I–V) based on the dynamics of CD20 expression during treatment (*lower panel*). Groups II and III correspond to cases with therapy-induced CD20 expression and/or strong upregulation. (**D**) Frequency of CD20^+^ blasts at diagnosis and at MRD Day15, stratified by genetic subtypes (*ETV6::RUNX1*, hyperdiploidy, *IgH::DUX4*, other B-ALL). Median of CD20 expression and percentage of positive patients (>20%) were indicated.

### Targeting CD20 with immunotherapy impairs MRD cells in PDX models

The above results indicate a phenotypic plasticity of blasts upon treatment, that could serve for the identification of novel vulnerabilities and be used for direct therapeutic interventions. Therefore, we postulated that the CD20 up-regulation in MRD cells can be exploited for anti-CD20 targeted therapy. Leukemic blasts from relapse HB#012 patient were transplanted and expanded into NSG mice (**Figures 6A**, **B**). 10 weeks after transplantation, highly engrafted mice (**Figure 6B**) were randomly selected for treatment or not (vehicle) with CHEMO for 3 weeks. Moreover, the anti-CD20 monoclonal antibody Obinutuzumab (OBINU) was added or not (CTL) in mice the last week of the treatment with CHEMO (**Figure 6A**). Thus, we confirmed that CHEMO efficiently decreased the proportion and the absolute number of leukemic blasts in the BM and the spleen of relapse PDXs (**Figures 6C-E**), associated with a massive reduction of the tumor load in the spleen (**Figures 6E**, **F**). However, CHEMO spared persisting MRD cells (**Figures 6C-E**) that have clearly up-regulated CD20 (**Figures 6G**, **H**), as expected. Strikingly, the combination of OBINU during the last week of CHEMO drastically impaired MRD cells (**Figures 6C-E**), closely eradicating CD20^+^ cells in the BM and the spleen of recipient mice (**Figures 6G-I**). Finally, we repeated this *in vivo* experimental approach with leukemic blasts from diagnosis HB#012 patient (**Figure S5A**) and demonstrated a clear synergistic antileukemic effect of OBINU and CHEMO cells in PDX models (**Figures S5B-F**). Collectively, our results demonstrate that glucocorticoid treatment of B-ALL cells upregulates CD20 expression, creating a synergistic vulnerability and a potent antileukemic effect on MRD cells with anti-CD20 monoclonal antibody in preclinical model.

**Figure 6.**
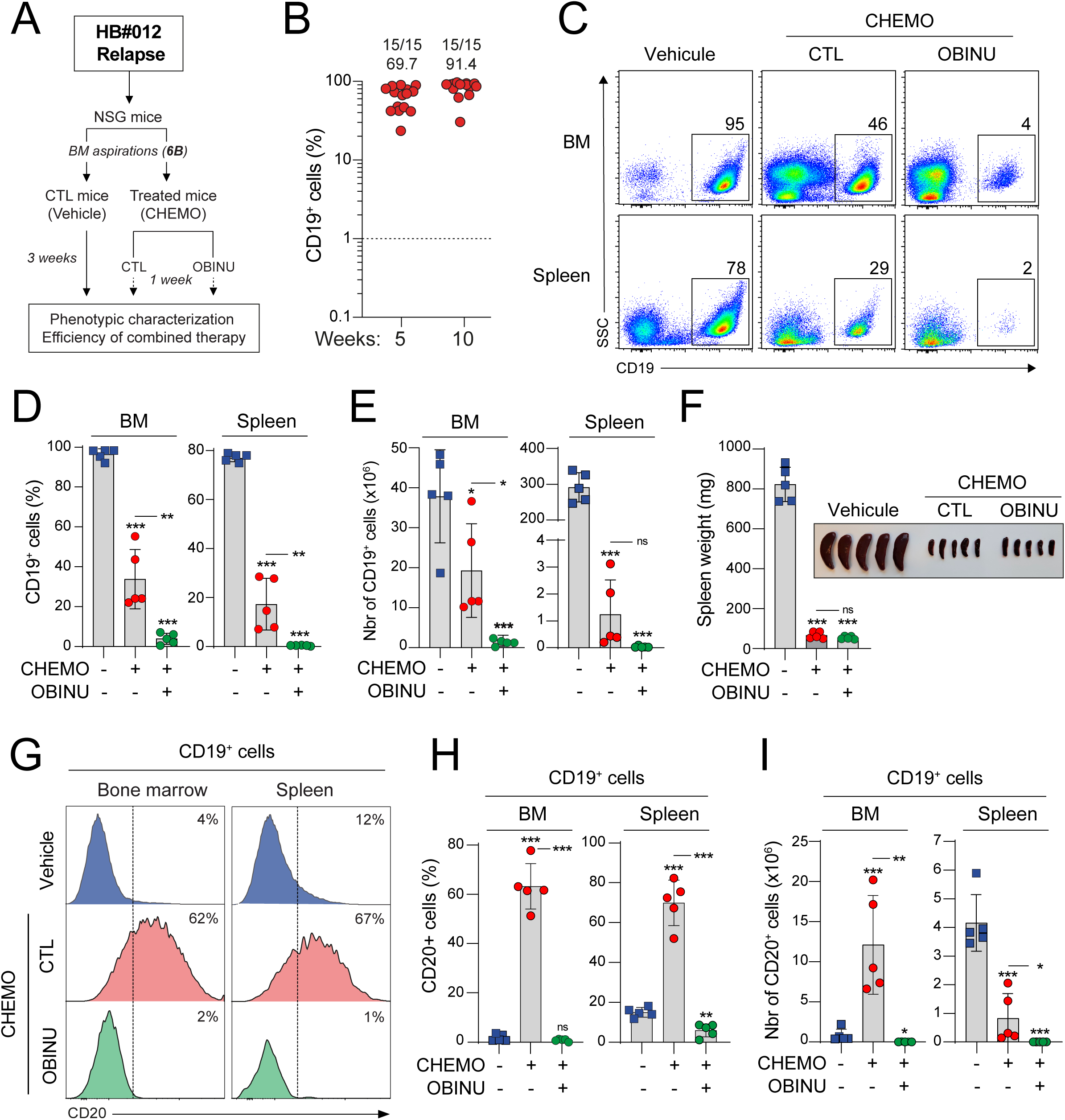
(**A-B**) Experimental procedure to evaluate the efficiency of the anti-CD20 monoclonal antibody Obinutuzumab in targeting MRD cells (A). Leukemic blasts from relapse HB#012 patient were transplanted into 15 NSG mice. After expansion (B), engrafted mice were randomly selected for treatment (CHEMO; n=10) or not (vehicle; n=5) with a chemotherapeutic cocktail (DEXA+VCR) during 3 weeks. Obinutuzumab (OBINU, n=5) was added twice or not (CTL, n=5) in mice the last week of the treatment with CHEMO (A). (**C-E**) Representative FACS analysis (C) and quantification (D) of the proportion of human CD19^+^ leukemic cells in the BM and the spleen after the treatment. Absolute numbers of CD19^+^ B-ALL blasts was calculated (E). (**F**) Spleen weight of each engrafted mouse was measured after treatment and picture of the representative spleens was shown. (**G-I**) Representative FACS analysis (G) and quantification (H) of CD20 expression in CD19⁺ B-ALL blasts from the BM and the spleen after the treatment. Absolute numbers of CD20^+^ B-ALL blasts was calculated (I).

## Discussion

By using PDX models of pediatric B-ALL, the present study uncovered a biological mechanism that occurs in drug-resistant cells after treatment with chemotherapies. We demonstrated that chemotherapies, in particular glucocorticoid treatment of B-ALLs, induce CD20 upregulation. Three notable features emerge from this work. First, glucocorticoid-triggered upregulation of CD20 occurs in the absence of detectable known mutations, indicating that CD20 activation may not be the result of the clonal evolution/selection within the tumor. Second, CD20 is consistently observed as an upregulated marker in MRD cells derived from PDXs and patients, irrespective of the oncogenic subtypes. Third, this mechanism provides a window of therapeutic opportunity for killing drug-resistant cells with anti-CD20 monoclonal antibodies.

Cell-cycle restriction is an important mechanism of therapeutic resistance, particularly for therapies targeting dividing leukemic cells. This notion is well exemplified at the clinical level in chronic myeloid leukemia (CML) where the absence of total cure, even after treatment with tyrosine kinase inhibitors, is due to the inability to eradicate quiescent residual clones (35–37). Cell-quiescence is also described as a crucial mechanism of drug-resistance in both human and murine B-ALL models (8–10). Herein, we explored the phenotypic and molecular heterogeneity of bulk cells at single cell resolution and uncovered a unique distinct MRD-like population exhibiting a mature immunophenotype CD45^high^CD34^low^CD20^+^ as well as a cell-quiescence signature. Notwithstanding the fact that drug resistance was focused in CD20^+^ quiescent cells, this observation raises at least three open questions: (i) whether cell-quiescence is a critical property to escape treatment and prime leukemia re-initiation, (ii) whether resistance is linked to a minor population of quiescent blasts within the tumor that are more differentiated along the B-cell lineage and (iii) whether cell-quiescence, drug resistance and leukemia initiation are plastic and reversible stem cell-like functions. Our results clearly demonstrate that CD20 is a plastic marker, as its expression is influenced by steroid exposure, and as it appears after the short-term engraftment of blasts that were initially CD20-negative. Importantly, our results show that this phenotypic plasticity is related neither to genetic heterogeneity nor to clonal selection and/or evolution processes of MRD cells. These observations therefore challenge the traditional stem cell hierarchy in leukemia and raise the question about whether surface marker plasticity is linked to functional plasticity within the tumor bulk. Supporting this idea, it has been shown both in B-ALL and T-ALL that cell-surface immunophenotype, dormancy and chemoresistance are influenced by BM niches, highlighting the cellular and functional adaptability of leukemic cells to their *in vivo* microenvironment (10–12).

Thus, our findings reveal that CD20 is not a fixed lineage marker, but a dynamically regulated molecule induced by glucocorticoid treatment. Although our study did not directly investigate the specific mechanism of CD20 modulation, we hypothesize that steroid hormones from induction therapy, as potent regulators of gene transcription (38), are central to this process. This antigen modulation is not restricted to CD20, as we also observed the down-regulation of CD34 and up-regulation of CD45 in MRD cells from our PDX models, as well as from patients. Indeed, immunophenotypic change affecting MRD markers is an important feature of leukemic cells upon treatment and is described in a large proportion of pediatric B-ALL patients (39–41). These observations raise important concerns for current MRD monitoring strategies. They highlight the limitations of static MRD panels and, consequently, support the need for dynamic gating strategies that take into account therapy-driven phenotypic plasticity. While the prognostic value of CD20 expression in pediatric B-ALL is debated in the literature (42–46), likely due to differences in patient cohort sizes and treatment protocols, the qualitative and quantitative up-regulation of CD20 after steroid exposure is a documented phenomenon (45,47,48). Herein, our study not only brings evidence that CD20 expression is modulated by chemotherapy, but also shows that modeling MRD using B-ALL PDX models accurately reflects the clinical situation in patients. Despite this obvious antigen modulation following treatment, our data revealed, nevertheless, a small and quiescent population with a mature CD34^low^CD20^+^ immunophenotype within the untreated tumor bulk of diagnosis PDXs. It is thus possible that more mature blasts, characterized by low CD34 expression and high CD20 expression, were selected by the treatment. Underrepresented at diagnosis, these cells could constitute the majority of MRD cells after treatment, thereby mimicking antigen modulation. This notion is especially supported by the demonstration that CD34^+^ leukemic cells from B-ALL patients are more sensitive and more rapidly depleted during chemotherapy than CD34^-^ cells (49). To precisely determine the contribution of antigenic modulation and/or the selection process during treatment, further studies are required. These could include cell tracking of CD20-positive and negative cell populations, followed by functional experiments.

Our observations also suggest that CD20 may functionally contribute to leukemic persistence. A plausible mechanism involves its role in supporting B-cell receptor (BCR) signaling, a pathway implicated in survival and therapy resistance. Indeed, several studies indicate that CD20 supports BCR signaling functions (50–53) and enhanced (pre-)BCR pathway activity has been associated with glucocorticoid resistance and poor clinical outcomes in B-ALL (54,55). Herein, our transcriptomic analyses revealed that CD20⁺ MRD cells exhibit strong enrichment for the (pre-)BCR signaling. This association is supported at the functional level showing that CD20^+^ cells robustly respond to (pre-)BCR stimulation through enhanced phosphorylation of PLCγ2. These findings support a model in which CD20 upregulation enhances the pro-survival and resistance of MRD cells by reinforcing the (pre-)BCR signaling. Importantly, B-ALL cells exhibiting a tonic (pre-)BCR signaling are sensitive to its downstream effectors SYK, SRC, BTK and PI3Kδ inhibition (51,56–59). Thus, loss-of-function approaches, as well as treatment with inhibitors of tonic (pre-)BCR such as Ibrutinib or Idelalisib would be of interest to fully explore the functions and the cross-talk between CD20 and BCR in MRD cells of B-ALL.

While CD20 monoclonal antibodies have shown clinical benefit in adult B-ALL (24), these therapies are not currently incorporated in pediatric B-ALL regiments. Here, our data clearly demonstrate that CD20 is robustly upregulated in MRD cells from PDXs and from about half of patients regardless of their oncogenic subtype, even those with low or undetectable levels at diagnosis. This dynamic regulation redefines CD20 as a transient and treatment-dependent vulnerability, and creates an actionable therapeutic opportunity during which anti-CD20 immunotherapy could be applied to eliminate drug-tolerant residual cells. Indeed, we demonstrated using our PDX models that combining induction chemotherapies with the Obinutuzumab significantly reduces MRD cells, providing a strong rationale for using this strategy to eliminate drug-tolerant leukemic cells. Thus, we propose that a subset of pediatric B-ALL patients could be stratified according to the ectopic activation of CD20 during induction chemotherapy, and could benefit from additional treatment with Obinutuzumab to reduce relapse risk.

## Materials and Methods

### Human B-ALL samples

Pediatric B-ALL samples were obtained from the Toulouse University Hospital (TUH) (Toulouse, France) and adult B-ALL samples from the Saint-Louis Hospital (Paris, France) after signed written informed consent for research use in accordance with the Declaration of Helsinki and stored at the HIMIP collection (BB-0033-00060) and at the BiRTH collection (“Biobanque pour la Recherche Translationnelle en Hématologie”, Paris). According to the French law, HIMIP and BiRTH biobank collections have been declared to the Ministry of Higher Education and Research (DC 2008-307 and DC 2009-929, respectively). We obtained a transfer agreement for research applications (AC 2008-129) after approbation by our institutional review board and ethics committee (Comité de Protection des Personnes Sud-Oust et Outremer II). Biological annotations of the samples have been declared to the CNIL (Comité National Informatique et Libertés, i.e., Data Processing and Liberties National Committee). Patients characteristics are provided in **Table S1**.

### Patient-Derived Xenografted Models and *in vivo* treatment with chemotherapy

Immunodeficient NOD.*Cg-Prkdc^scid^Il2rg^tmWjl^*(NSG) mice were produced at the Génotoul-Anexplo platform at Toulouse, France, using breeders obtained from Charles River Laboratories. Mice were housed in sterile conditions using high-efficiency particulate arrestance filtered microisolators and fed with irradiated food and sterile water. Mice (6–9 weeks old) were sub-lethally treated with 30 mg/kg of busulfan 24 h before injection of leukemic cells. Leukemic blasts from diagnosis or relapse B-ALL patients were thawed at room temperature, washed twice in PBS and transplanted by intravenous injection (2×10^5^ cells per mouse). Leukemia engraftment was monitored at the indicated time points by bone marrow aspirations followed by flow cytometry using antibodies against human CD45, CD19 and CD20 markers (BD Biosciences). Mice displaying more that 70% of human leukemic engraftment in the bone marrow were randomly assigned to control or treatment groups. For combination therapy, a chemotherapeutic cocktail (CHEMO) composed of dexamethasone (DEXA, 10 mg/kg) and vincristine (VCR, 0.5 mg/kg) was administrated weekly for 3 weeks *via* intraperitoneal injection. A daily administration of 10 mg/kg DEXA was supplemented during the period of treatment. The anti-CD20 monoclonal antibody Obinutuzumab (OBINU, 20 mg/kg) was intraperitoneally administrated twice (days 15 and 19) the last week of CHEMO treatment. Mice were euthanized and analysed after the treatment.

### FACS analysis, antibodies and cell sorting

Cells freshly harvested were treated with ACK buffer to lyse red blood cells before staining. Single cell suspensions from human B-ALL cells were prepared in Iscove’s Modified Dulbecco’s Medium (IMDM; Gibco) supplemented with 2% Fetal Bovine Serum (FBS) (Stemcell technologies). Immunostainings were performed using antibodies for flow cytometry obtained from Pharmingen (BD Biosciences), are listed in **Table S5**. For surface staining, cells were incubated with the antibodies for 20 minutes in IMDM 2% FBS at 4°C and washed twice with PBS1X before analysis. FACS analysis was performed on a Fortessa cytometer (BD Biosciences) using FlowJo (BD Biosciences) software and cell sorting was performed on a FACS Melody cell sorter (BD Biosciences).

### UMAP analysis

UMAP (Uniform Manifold Approximation and Projection) analysis was performed using FlowJo software using a multiparametric staining by FACS integrating the seven markers. Samples were pre-gated on single cells of the population of interest. Then, informatic cleaning and normalization were performed using FlowAI package (V2.1) and CytoNorm package (V1.0) for each sample and the same number of cells were then concatenated. Arc sin transformation was performed manually for each marker to discriminate the different B-cell populations. The dimension reduction algorithm UMAP (umap-learn Python package v2.4.0) was run using Euclidian distance with 15 nearest neighbors and 0.5 distance parameters.

### (Pre-)BCR activation and phosphoflow cytometry

To assess the functional integrity of the (pre-)B cell receptor signaling pathway in leukemic cells, we performed an *ex vivo* stimulation assay based on hydrogen peroxide (H₂O₂)–induced activation, as previously described (8). This approach allows the detection of proximal BCR signaling events by measuring phosphorylation of key adaptor proteins such PLCγ2 by phospho-flow cytometry. (Pre-)BCR activation was assessed by phospho-flow cytometry on thawed leukemic cells from pediatric B-ALL patients. Viable CD20⁺ and CD20- cells were sorted by FACS prior to stimulation. Cells were then incubated for 5 minutes at 37 °C in RPMI 1640 supplemented with 2% FBS, either in the presence or absence of 1 mM hydrogen peroxide (H₂O₂) (Sigma-Aldrich), to stimulate (pre-)BCR signaling. Following stimulation, cells were fixed and permeabilized using Cytofix/Cytoperm Plus buffer (BD Biosciences) for 45 minutes at 4 °C. Cells were then washed twice with Perm/Wash buffer (BD Biosciences). Intracellular staining was performed using fluorochrome-conjugated antibodies against phospho-PLCγ2 (Invitrogen, 12-9866-42), along with viability and lineage markers. Samples were acquired on a BD LSRFortessa flow cytometer, and data were analyzed using FlowJo software.

### *In vitro* co-culture assays and dexamethasone treatment

Human B-ALL cells from PDXs were washed, filtered through a 70 µm cell strainer, and resuspended in Iscove’s Modified Dulbecco’s Medium (IMDM; Gibco) supplemented with 5% fetal bovine serum (FBS; Stemcell Technologies), 0.05 mM β-mercaptoethanol (Sigma-Aldrich), 2 mM L-glutamine (Invitrogen), 100 U/mL penicillin, 100 U/mL streptomycin, and the following recombinant cytokines from PeproTech: IL-7 (5 ng/mL), FLT3 ligand (10 ng/mL), and stem cell factor (SCF; 10 ng/mL). Cells were co-cultured onto a monolayer of MS-5 stromal cells (purchased from DSMZ) previously plated to adhere overnight. Co-cultures were maintained at 37 °C and 5% CO₂ and treated for 48 hours, either with vehicle (control), with a dose-response range of DEXA, or with a single dose. At the end of the culture period, non-adherent cells were carefully collected, counted, and washed. Surface staining was performed using fluorochrome-conjugated antibodies, including anti-CD20, and Zombie viability dye (BioLegend) was used for dead cell exclusion.

### Mutational screening and next generation sequencing (NGS) of targeted gene regions

DNA extraction was performed using a AllPrep® DNA/RNA micro-Kit (#80284) according to manufacturer’s instruction (Qiagen). An extended DNA sequencing was performed on leukemic blasts from HB#012 patient at diagnosis and relapse, analyzing the complete coding region of 95 genes (**Table S6**). Libraries were prepared using SureSelect custom panel (Agilent) according to manufacturer instructions and sequenced using an Illumina NextSeq500. Alignment was performed using BWA aligner and variant calling was performed using HaplotypeCaller, Mutect2 and Surecall (Agilent) variant callers. A sensitivity threshold of 1% variant allele frequency was used for mutation reporting and manual curation. The recurrence of *KRAS* mutation, as well as IG/TCR gene rearrangements, were screened by targeted NGS in leukemic cells from the BM of HB#012 diagnosis and relapse PDXs treated or not with CHEMO. Targets and primers are listed in **Table S7**.

### Bulk RNA-sequencing

Human B-ALL cells were purified from PDXs, resuspended in RLT buffer and the RNA was isolated with the RNeasy plus micro kit (Qiagen). Smart-seqv4 libraries were prepared as previously described (60) using the Takara SMART-Seq v4 full-length transcriptiome analysis kit according to the manufacture’s guidelines. Paired-end sequencing was performed on an Illumina NextSeq 500 using 2 x 75 bp reads. Low quality reads were trimmed and adaptor sequence was removed using trimmomatic. Short reads were then mapped to GRCh38.p14 genome using HISAT2. Low quality mapping and duplicated reads were removed and the remaining reads were count using FeatureCounts with default paired-end parameter and regularized log-transformed (rlog) values were calculated by DESeq2. Differentially expressed genes (RNA-seq signal) have been analysed using the online iDEP2.01 software (http://bioinformatics.sdstate.edu/idep/).

### Single-cell RNA-sequencing

Human B-ALL cells were purified from treated and untreated PDXs and single-cell transcriptomic analyses were performed using the Chromium Single Cell Gene Expression FLEX platform (10x Genomics), following the manufacturer’s protocol. Fixed cell suspensions were prepared using the Chromium Next GEM Single Cell Fixed RNA Sample Preparation Kit from B-ALL PDX-derived samples, and approximately 50,000 cells per sample were loaded for each reaction. Library preparation, barcoding, and cDNA synthesis were carried out using the Chromium Fixed RNA Profiling Reagent kits, optimized for fixed cells. Libraries were sequenced on an Illumina platform (NovaSeq 6000) to ensure sufficient read depth per cell. Raw sequencing data were processed using the Cell Ranger pipeline (10x Genomics) for alignment, barcode processing, and UMI counting. The resulting gene expression matrices were analyzed with the Seurat R package (version 5.2.1) for quality control, normalization, dimensionality reduction, and clustering. Principal component analysis (PCA) and Uniform Manifold Approximation and Projection (UMAP) were used for visualization of cell populations. Cluster-specific marker genes were identified through differential gene expression analysis. Gene signature analysis was performed using Ucell R package with Seurat. Gene sets were curated from two sources: (i) previously published and peer-reviewed studies relevant to leukemia biology and treatment resistance, and (ii) publicly available gene sets from the Molecular Signatures Database (MSigDB) (https://www.gsea-msigdb.org), including hallmark pathways and Gene Ontology categories. In addition, a curated list of genes encoding surface proteins was extracted from The Cancer Surfaceome Atlas (25).

### Immunophenotyping of MRD cells in pediatric B-ALL patients

Immunophenotypic analyses were performed on bone marrow aspirates collected at diagnosis (day 0) and at day 15 of induction therapy, in patients treated in the ALLTogether-1 protocol (NCT04307576). A total of 340 pediatric B-ALL patients were screened, and 45 patients with MRD >0.3% at day 15 were selected for extended CD20 analysis. Flow cytometry was conducted on fresh or cryopreserved mononuclear cells using panels that systematically included CD19, CD20, CD45, and CD34. At the day 15, additional markers such as CD10, CD38, and CD73 were routinely used to refine blast gating and assess phenotypic changes. Other antibodies were added depending on the immunophenotype at diagnosis or specific clinical features. All flow cytometry analyses were performed by experienced clinical cytometrists specialized in hematological malignancies, within accredited hospital diagnostic laboratories. Patient information’s and flow cytometry data are detailed in **Table S8**. This study was approved and supported by the ALLTogether research consortium and used data and/or samples from patients treated on one or more ALLTogether clinical trials. We thank all the patients, and their families, who participated in the research. We acknowledge the contribution of the clinicians, scientists and administrators who established the ALLTogether consortium and conducted the clinical trials.”

### Statistical analysis

Student’s *t* test was used for comparison of quantitative variable (\*\*\**p*<0.0005, \*\**p*<0.005, \**p*< 0.05), assuming normality and equal distribution of variance between the different groups analyzed.

## Data availability statement

Bulk and single cell RNA-seq data reported in this study are available at the Gene Expression Omnibus (GEO) repository under the accession number GSE309451. All other raw data generated in this study are available upon request from the corresponding author.

## Acknowledgments

We acknowledge Manon Farcé from the cell sorting facility of the Cancer Research Center of Toulouse (INSERM U1037) and the Anexplo/Genotoul platforms (UMS006) for technical assistances. This study was supported by institutional grants from the Institut National de la Santé et de la Recherche Médicale (INSERM, France), France, from the Centre National de la Recherche Scientifique (CNRS, France), the Institut National du Cancer, France (INCa-2020-096, France), the Agence Nationale de la Recherche (ANR-18-CE13-0002-01, France), the Fondation ARC pour la Recherche sur le Cancer (PJA-20181207977, France), the Fondation de France (WB-2023-49770, France) and the CARe graduate school. The team is labelled by the Ligue Contre le Cancer and supported by the associations “Laurette Fugain”, “111 des arts”, “Cassandra” and “Constance la petite guerrière astronaute”.

## Author contributions

J. Bigot designed the study, performed and analysed the experiments and wrote the manuscript. M. Bouttier, V. Fregona, C. Rouzier, M. Bayet, S. Hebrard, N. Prade and S. Lagarde conceived and performed experiments. F. Vergez, A. Baruchel, M. Strullu and E. Lainey provided the data from B-ALL patients. A. Quillet-Mary and L. Ysebaert provided the Obinutuzumab. E. Clappier generated and provided P80R B-ALL cells from PDXs. C. Didier, L. Largeaud, C. Broccardo, M. Pasquet and E. Delabesse reviewed the data and the manuscript. B. Gerby acquired the funding, performed and analysed experiments, conceived and supervised the project and wrote the manuscript. All authors contributed to the final draft.

## Supplementary figure legends

**Figure S1.** (**A-B**) Representative FACS analysis of size and granularity (*upper panel*) and of the proportion of human CD19^+^ leukemic cells (*lower panel*) in the BM and the spleen from both diagnosis and relapse HB#012 PDXs treated or not (vehicle) with CHEMO (A). Absolute numbers of CD19^+^ B-ALL blasts was calculated in the BM and the spleen of diagnosis and relapse PDXs (B). (**C-E**) Experimental procedure to explore the gene expression profile of chemo-resistant B-ALL cells (C). Leukemic blasts from the “*de novo*” B-ALL patient HB#010 were transplanted into NSG mice (n=10) and engrafted mice were randomly selected for treatment (CHEMO, n= 5) or not (vehicle, n=5) with a chemotherapeutic cocktail (DEXA+VCR) during 3 weeks, as previously described (8). Representative FACS analysis (D) and quantification (E) of the reconstitution (proportion of CD19^+^ B-ALL blasts) in the BM and the spleen after the treatment. (**F**) Volcano plot showing differentially expressed genes (RNA-seq signal) between vehicle (n=3) and CHEMO (n=3) leukemic HB#010 cells. Up-regulated (red dots) and down-regulated (blue dots) genes are selected for an expression difference of >3-fold (FDR<0.05). (**G-I**) GSEA of the up- (*left panel*) and down- (*right panel*) regulated genes from residual human B-ALL cells after chemotherapies (MRD signature) (10) between vehicle and CHEMO cells (G). GSEA of the molecular signatures “TNFα signaling”, “Inflammatory response”, “Hypoxia” (H) and “E2F targets” (I) between vehicle and CHEMO cells. The normalized enrichment score (NES) and the false discovery rate (FDR) are indicated.

**Figure S2.** (**A**) Gene signatures of cell cycle phases (G0/1, S and G2+M) (*upper panels*) and normalized expression levels of *MKi67* (*lower panels*) were shown on the UMAP representation of the scRNA-seq data from diagnosis and relapse HB#012 PDXs treated or not with CHEMO. (**B**) Venn diagram indicating the overlap of the list of up-regulated genes encoding surface makers in MRD cells from diagnosis and relapse HB#012 PDXs (scRNA-seq) and the list of up-regulated genes in MRD cells from HB#010 PDXs (bulk RNA-seq). The list of overlapping genes is indicated. (**C-E**) Enrichment scores of primary MRD signature (10) (C, *left panel*), of cell-quiescence signature (29) (C, *right panel*), as well as of BCR signaling pathway (D, GO and KEGG databases) were calculated and displayed across the clusters of the scRNA-seq data from diagnosis untreated HB#012 PDXs. Cluster 10 was colored in purple. Enrichment scores of these three signatures were also analyzed between *MS4A1* positive and negative cells from the scRNA-seq data (E). (**F**) Schematic representation of the human B-cell development. Representation of the different B-cell differentiation stages (CLP, pre-pro-B, pro-B, pre-BI, pre-BII, immature-B and mature-B) and expression patterns of the main markers characterizing each subset. (**G**) Visualization UMAP of the expression level of CD10 and CD38 on human CD19⁺ B-ALL blasts from the BM of diagnosis and relapse HB#012 PDXs treated (CHEMO) or not (Vehicle).

**Figure S3.** (**A**) KRAS^G12R^ mutation and TCRδV2-D3 rearrangement found in HB#012 patient at diagnosis and relapse were screened for recurrence by targeted next-generation sequencing (NGS) in leukemic cells from the BM of PDXs treated or not with CHEMO. Numbers represent the variant allele frequency (VAF) of the KRAS mutation and of the TCRδ rearrangement. (**B**) KRAS^G12R^ mutation and TCRδV2-D3 rearrangement were analyzed on purified CD20^+^ and CD20^-^ leukemic cells from diagnosis PDXs. (**C**) Quantification of CD20 expression in CD19⁺ B-ALL blasts from the BM and the spleen of from HB#010 PDXs treated or not with CHEMO. (**D**) Spleen weight of diagnosis and relapse HB#020 PDXs treated or not with CHEMO. (**E-F**) Human B-ALL reconstitution (% of CD19^+^ B-ALL blasts) was monitored by FACS in the BM and the spleen of diagnosis and relapse HB#020 PDXs treated or not with CHEMO (E) and representative FACS analysis of CD20 expression in CD19⁺ B-ALL blasts was shown (F). (**G**) Representative FACS analysis (*left panel*) and quantification (*right panel*) of phosphorylated PLCγ2 (pPLCγ2) in purified CD20⁺ and CD20⁻ leukemic cells from relapse HB#012 PDXs after *ex vivo* BCR stimulation using hydrogen peroxide (+H₂O₂). Unstimulated cells (-H₂O₂) served as controls. (**H**) CD20 expression on leukemic cells from relapse HB#012 PDXs after *in vitro* dose-response of DEXA. (**I**) Proportion of CD20⁺ B-ALL blasts (CD19⁺) (*left panel*) and absolute number of CD19^+^ B-ALL blasts (*right panel*) was measured after the *in vitro* dose-response of DEXA on purified CD20^+^ (blue) and CD20^-^ (red) leukemic cells from relapse HB#012 PDXs.

**Figure S4.** (**A**) Representative FACS analysis of CD20 expression on leukemic blasts at diagnosis and at MRD (day 15) for each patient group (I–V). (**B**) Heatmap showing the proportion of CD20^+^ cells at diagnosis and at day 15 across the different genetic subtypes.

Figure S5. (**A**) Leukemic blasts from diagnosis HB#012 patient were transplanted into 7 NSG mice and treated with CHEMO during 3 weeks. Obinutuzumab (OBINU, n=3) was added twice or not (CTL, n=4) in mice the last week of the treatment with CHEMO. (**B-D**) Representative FACS analysis (B) and quantification (C) of the proportion of human CD19^+^ leukemic cells in the BM and the spleen after the treatment. Absolute numbers of CD19^+^ B-ALL blasts was calculated (D). (**E-F**) Quantification (E) and absolute number (F) of CD20^+^ cells within CD19^+^ B-ALL blasts.

