## Supplementary figures and images for "Targeting glucocorticoid-induced CD20 activation in preclinical models of B-ALL"

### Supplemental S1-S5

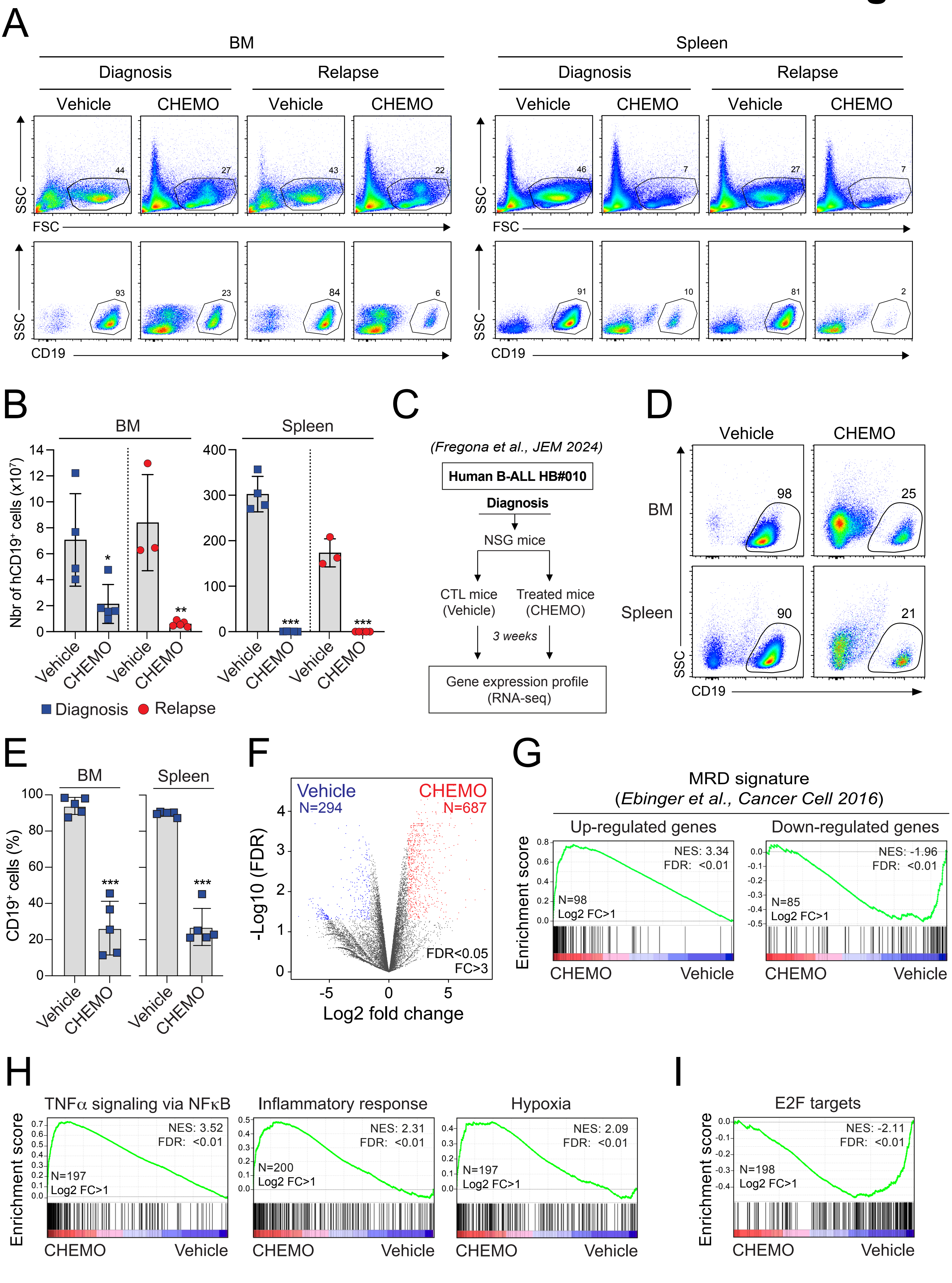

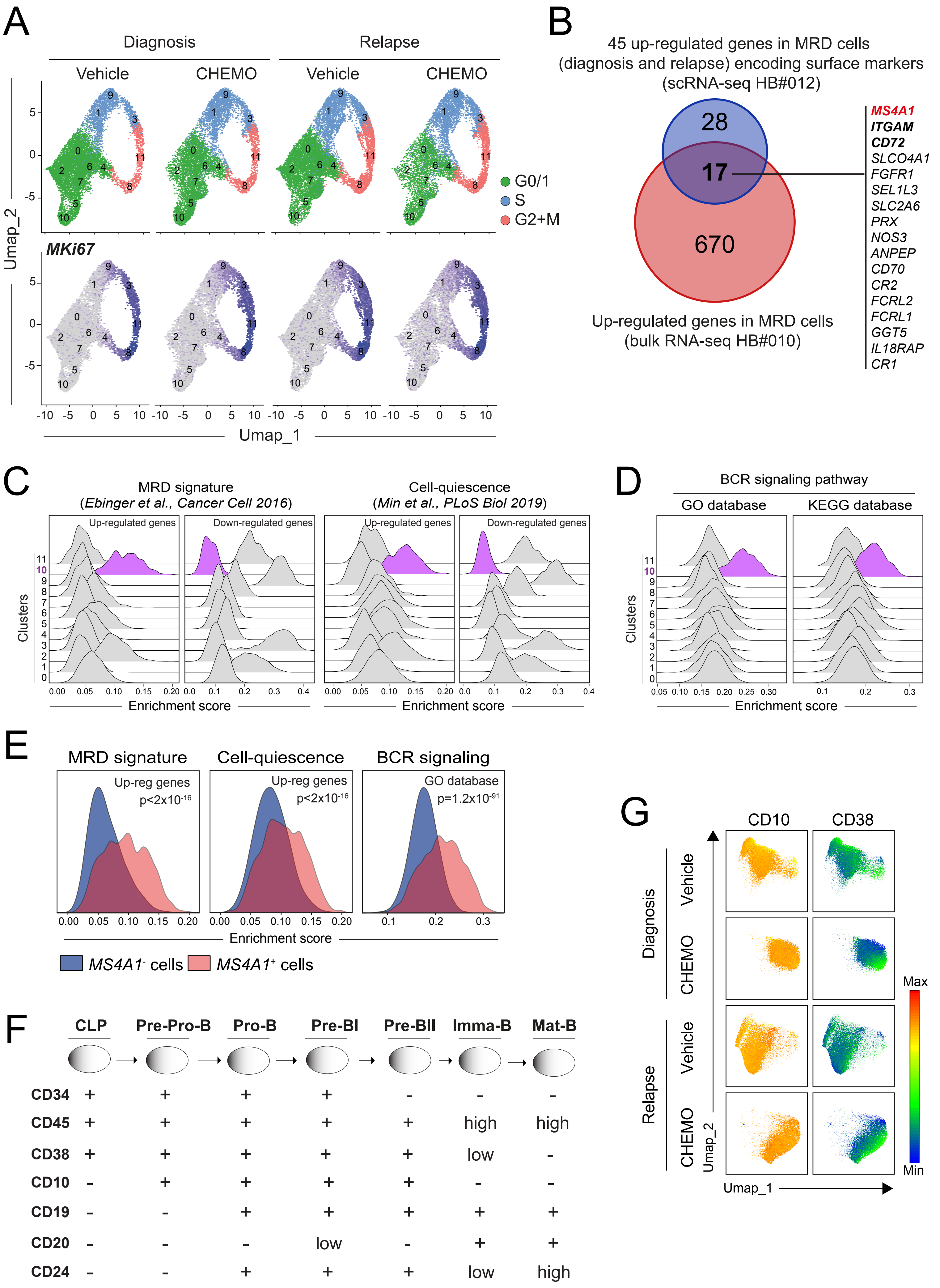

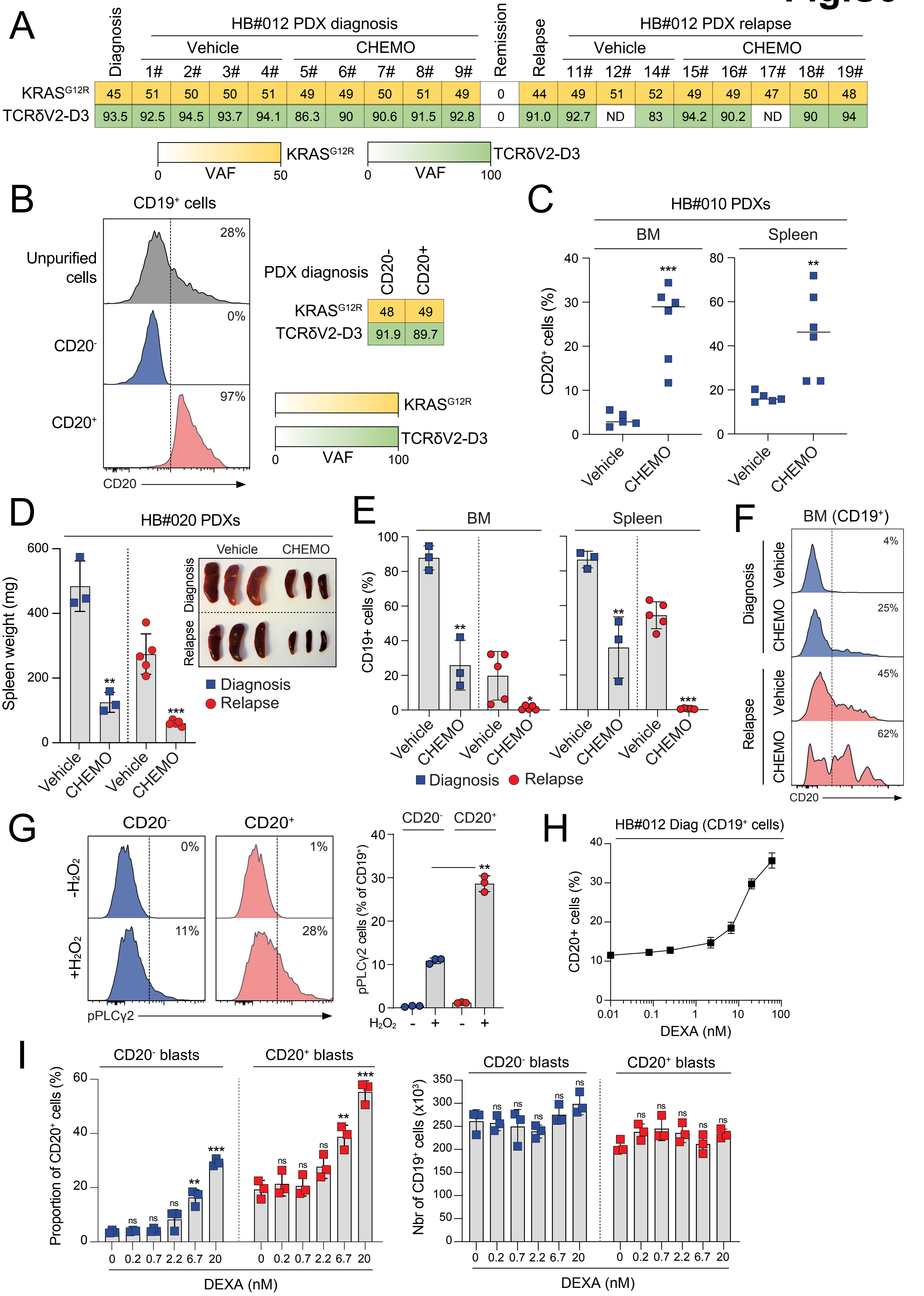

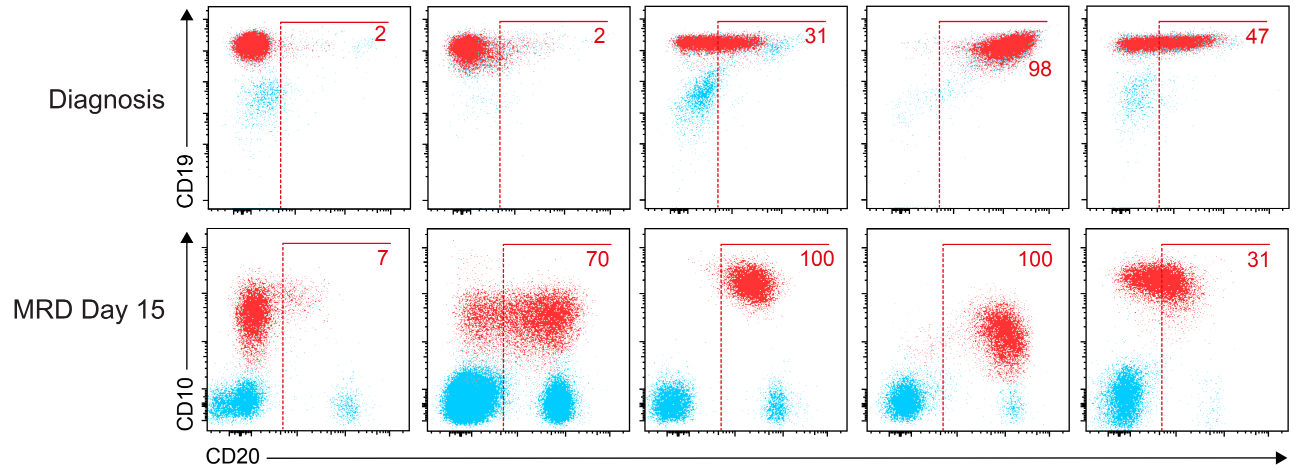

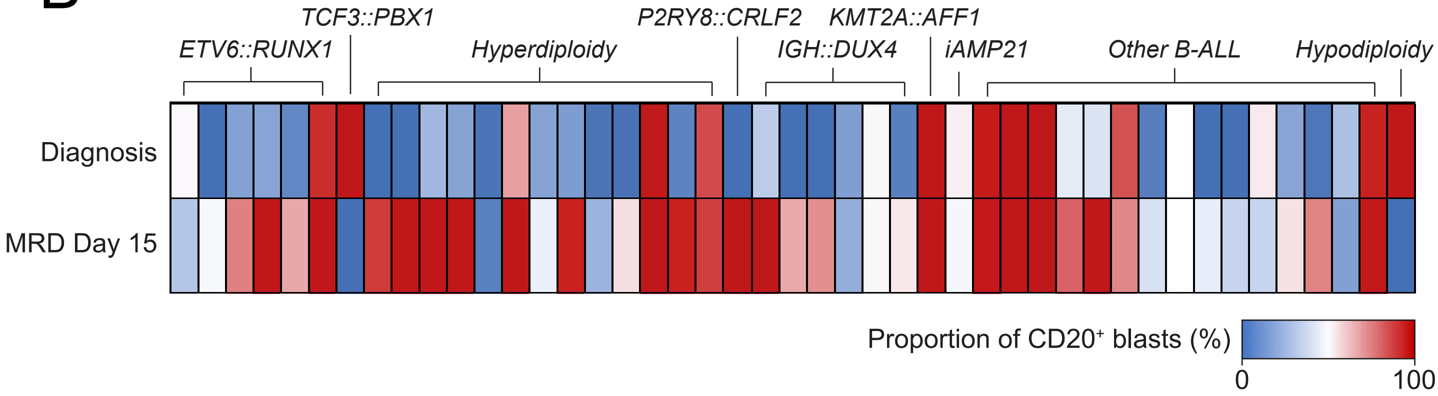

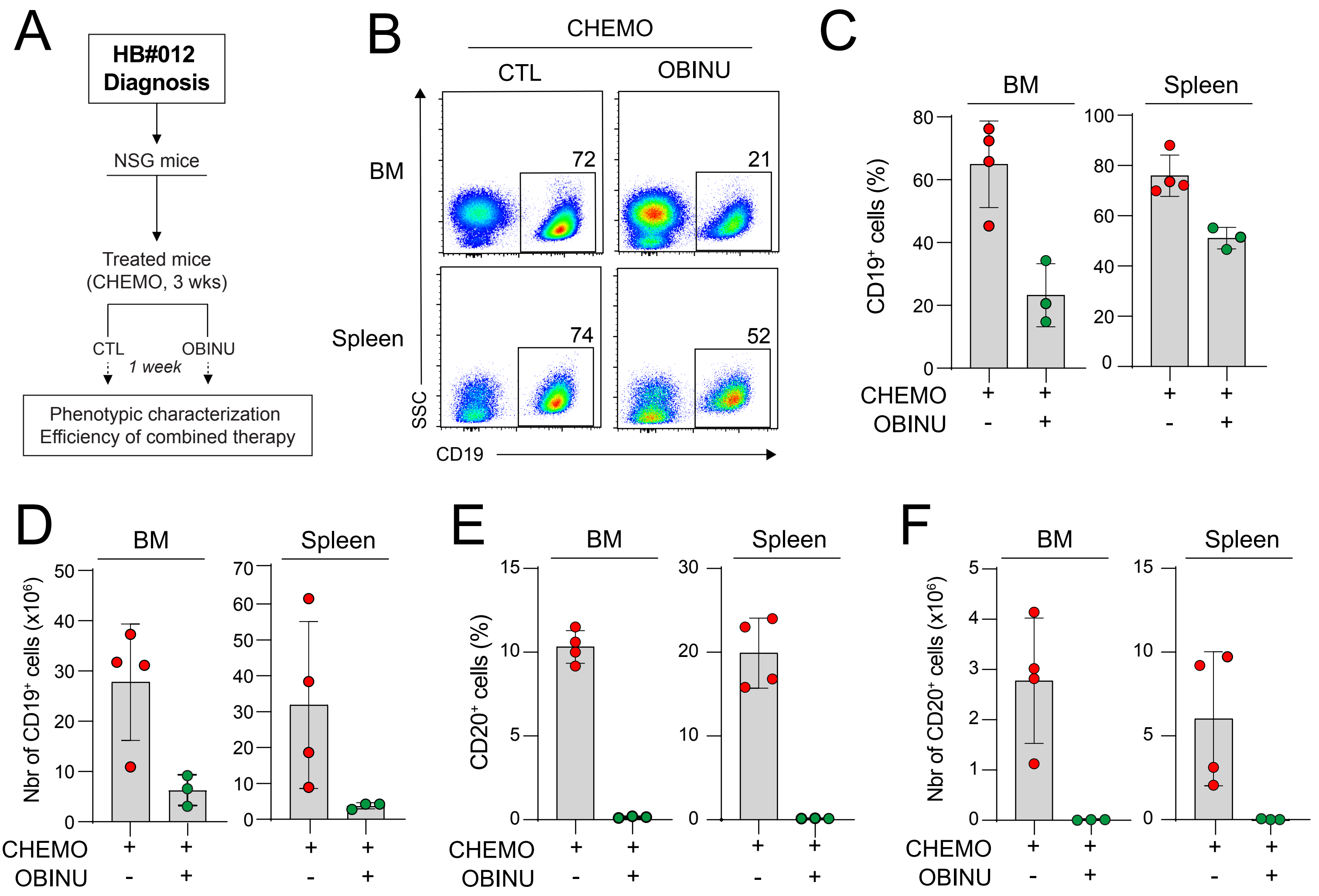
